# Autonomous multicolor bioluminescence imaging in bacteria, mammalian, and plant hosts

**DOI:** 10.1101/2024.04.28.591567

**Authors:** Subhan Hadi Kusuma, Mitsuru Hattori, Takeharu Nagai

**Affiliations:** Graduate School of Frontier Biosciences, Osaka University, Suita, Osaka 565-0871, Japan; SANKEN (The Institute of Scientific and Industrial Research), Osaka University, Ibaraki, Osaka 567-0047, Japan; OTRI, Osaka University, 1-1 Yamadaoka, Suita, Osaka 565-0871, Japan; Research Institute for Electronic Science, Hokkaido University

**Keywords:** Bioluminescence, Bacterial Luciferase, Biosensor

## Abstract

Bioluminescence imaging has become a valuable tool in biological research, offering several advantages over fluorescence-based techniques, including the absence of phototoxicity and photobleaching, along with a higher signal-to-noise ratio. Common bioluminescence imaging methods often require the addition of an external chemical substrate (luciferin), which can result in a decrease in luminescence intensity over time and limit prolonged observations. Since the bacterial bioluminescence system is genetically encoded for luciferase-luciferin production, it enables autonomous bioluminescence (auto-bioluminescence) imaging. However, its application to multiple reporters is restricted due to a limited range of color variants. Here, we report five colors auto-bioluminescence system named Nano-lanternX (NLX), which can be expressed in bacterial, mammalian, and plant hosts, thereby enabling auto-bioluminescence in various living organisms. We have also expanded the applications of the NLX system, such as multiplexed auto-bioluminescence imaging for gene expression, protein localization, and dynamics of biomolecules within living mammalian cells.

## Introduction

Bioluminescence is the production of light through an enzymatic reaction involving oxidation of a chemical substrate (luciferin) by an enzyme (luciferase) (1). The light produced by bioluminescent reactions is widely used to observe biological phenomena in living cells. To enable real-time imaging, continuous addition of luciferin is required. This can increase luciferin’s autooxidation (2), lowering the signal-to-noise ratio and complicating long-term imaging. To overcome this issue, efforts have been made to produce luciferin-related genes to create autonomous bioluminescence (auto-bioluminescence) systems. Bacterial luciferases (Lux) and fungal luciferases (Luz) are examples of such systems, but Luz is temperature-sensitive and has poor solubility (3), making Lux a more preferred option for auto-bioluminescent probes due to its physicochemical properties.

Lux is a heterodimeric luciferase (LuxA and LuxB), which generates blue-green light (490 nm) by oxidizing reduced flavin mononucleotide (FMNH_2,_ produced by flavin reductase/Frp) and long-chain fatty aldehyde (RCHO, produced by LuxCDE enzymes) to flavin mononucleotide (FMN) and to carboxylic acid (RCOOH), respectively(4). However, when observing multiple biological events, Lux-based probes cannot be used because of the lack of distinct color variants compared to commonly modified luciferases (5, 6). Consequently, there is a considerable demand for Lux color variants to enable monitoring of diverse biological events. Previous color variants of Lux with a small emission shifted (4-16 nm) typically emerge as a result of mutations occurring in the active site of luciferase (7) or through protein-binding interactions with yellow fluorescent protein (YFP) (8, 9) and lumazine protein (LumP) (10, 11), but these affect Lux activity, resulting in reduced luminescence (7) and thermostability (9–11).

To circumvent these limitations, we adopted a bioluminescence resonance energy transfer (BRET) strategy inspired by Nano-lantern development (5, 6, 12), which successfully produced multicolor variants of luciferase without affecting its luminescence and thermostability. Here, we introduce the multicolor auto-bioluminescent Lux for multiplexed imaging, expressible in bacterial, mammalian, and plant hosts. These color variants allow the visualization of multiplex gene reporters and subcellular tags. We also developed peptide-based indicators based on multicolor Lux

## Results

### Development of multicolor Lux

To change Lux’s bioluminescence color, we adopted a BRET strategy using the yellow fluorescent protein (FP) Venus (13) as the initial BRET acceptor protein (Fig. 1*A*). The effects of BRET with Venus, including modifications, were verified as previously reported (14). Venus was fused to the N- or C-terminus of the active subunit of Lux (LuxA) to achieve higher BRET efficiency (*SI Appendix*, Fig. S1*A*). To compare the bioluminescence intensities of these fusion proteins without the influence of endogenous FMN levels, we measured them in purified proteins, not in cell systems, as previously reported (14). For bioluminescence measurements, purified Lux and Venus fusions were added to FMN, decanal, and BNAH (15). The intensities of these fusions were found to be lower than those of wild-type Lux (*SI Appendix*, Fig. S1*B*). We speculated that unstructured residues from the C-terminus of Venus might affect the activity or protein folding of LuxA. Consequently, deleting Venus’s C-terminus (VenusΔC10-LuxA) significantly improved the bioluminescence intensity compared to wild-type (*SI Appendix*, Fig. S1*B*). Referring to this structure, other FPs, including mTurquoise2 (16) (cyan FP), sfGFP (17) (green FP), mKOκ (18) (orange FP), and mScarlet-I (19) (red FP), were fused with LuxA to serve as BRET acceptors. In the development of the red variant, the selection of FP was carried out first because of the lower spectral overlap between Lux and common red FPs. mScarlet-I (19) was selected due to its higher BRET efficiency compared to CyOFP1 (20), mCherry-XL (21), and Scarlet (19) (*SI Appendix*, Fig. S2 and Table S1).

**Fig. 1.**
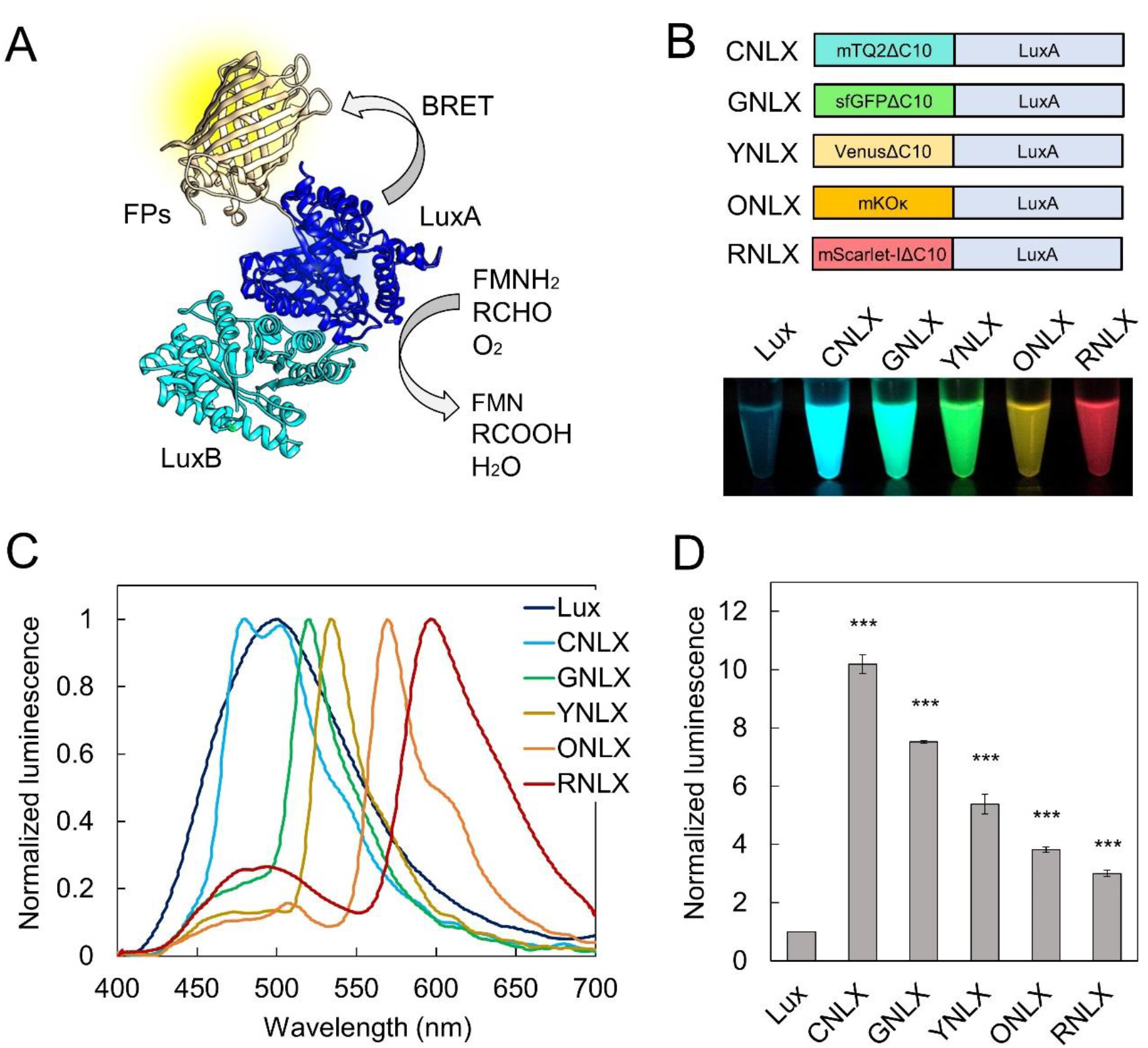
Development and characterization of NLX. (*A*) Structural model of NLX, consisting of heterodimeric Lux, LuxA (blue), LuxB (cyan), and FP (yellow). (*B*) Schematic of the NLX series (upper panel) and bioluminescence images of recombinant NLXs proteins (bottom). (*C*) Bioluminescence spectra of NLXs proteins. The bioluminescence intensities were normalized to each peak intensity. (*D*) Bioluminescence intensities of NLXs. Comparison of bioluminescence intensities of NLXs proteins and Lux. Data are mean ± s.d. *n* = 3, \*\*\**p* < 0.001.

We also attempted to fusing FPs to the N- or C-terminus of LuxB to assess their suitability for LuxB fusion (14) aiming to produce a multicolor Lux. Even then, the BRET efficiency in the LuxA fusion was higher than that in the LuxB fusion (*SI Appendix*, Fig. S3 and Table S1). Therefore, we designated these LuxA-based variants as NLX (**N**ano-**l**antern based on Lu**x** luciferase), consisting of cyan NLX (CNLX) (*λ*_max_^EM^=480 nm), green NLX (GNLX) (*λ*_max_^EM^=520 nm), yellow NLX (YNLX) (*λ*_max_^EM^=534 nm), orange NLX (ONLX) (*λ*_max_^EM^=569 nm), and red NLX (RNLX) (*λ*max^EM^=597 nm) (Figs. 1*B* and 1*C*). The bioluminescence intensities of the purified NLXs were higher than those of the original Lux, although the respective increases were different (Fig. 1*D*). The fusion of FP increased the luminescent quantum yield (QY), while the enzymatic parameters (*k*_cat_) of all NLXs were similar to those of wild-type Lux (*SI Appendix*, Figs. S4*A*-4*B* and Table S2). As reflected by the emission intensity, the QY of CNLX was six times higher than that of Lux (*SI Appendix*, Table S2).

### Autonomous bioluminescence imaging of NLXs

We next evaluated co-expression of NLXs with luciferin biosynthesis genes, such as *frp* and *luxCDE*, to promote multicolor auto-bioluminescence in various living organisms. We initially examined multicolor auto-bioluminescence in bacterial cell hosts. This involved transformation of *E. coli* with NLXs and the *luxCDE* operon, both driven by the T7 promoter. The bioluminescence emitted by recombinant *E. coli* expressing individual NLXs exhibited the expected colors, consistent with the bioluminescence observed in NLX-purified proteins (Fig. 2*A*). Additionally, we conducted multiplexed auto-bioluminescence imaging by mixing *E. coli* containing CNLX, YNLX, and RNLX plasmids (Fig. 2*B*). Recombinant *E. coli* demonstrated the capacity to be differentiated by color, using specific optical filters, and minimizing spectral noise with spectral unmixing algorithms (22). This study marks the first report of multicolor auto-bioluminescent bacteria that have a large spectral range from blue to red.

**Fig. 2.**
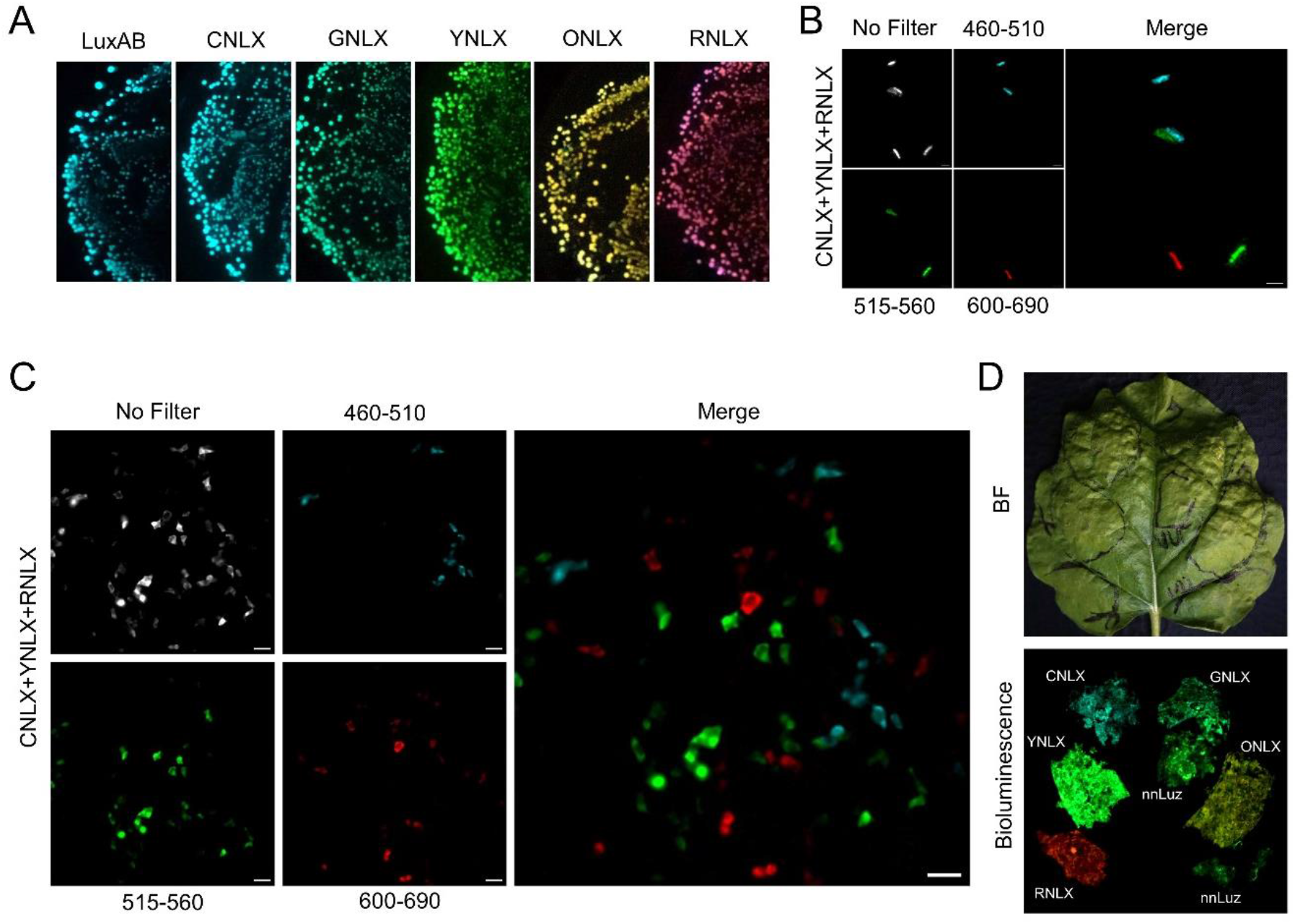
Multicolor auto-bioluminescence imaging with NLXs. (*A*) Auto-bioluminescence images of *E. coli* expressing all NLX variants. Bioluminescence images were acquired using a Sony α7s camera. (*B*) Auto-bioluminescence imaging of mixed *E. coli* expressing CNLX, YNLX, or RNLX. Scale bars, 2 μm: ×100 magnification. (*C*) Auto-bioluminescence imaging of mixed HEK293T cells expressing CNLX, YNLX, or RNLX. Scale bars, 50 μm: ×40 magnification. (*D*) Auto-bioluminescence of *Nicotiana benthamiana* leaves expressing NLXs genes and nnLuz. Bioluminescence images were acquired using a Sony α7s camera.

To evaluate NLXs’ performance in mammalian cells, they were introduced into the human codon-optimized *lux* operon (co *luxA*, co *luxB*, co *luxC*, co *luxD*, co *luxE*, and co *Frp*) (23), replacing *luxA* in NLXs with co *luxA*. Since the original co lux expression necessitated co-transfecting six plasmids for each gene, we sought to minimize the number of plasmids by utilizing the 2A peptide to link proteins (*SI Appendix*, Fig. S5*A*). co *luxD*, co *luxE*, and co *luxC* were constructed as a tricistronic unit using two 2A peptides (co *luxD*-*P2A*-co *luxE*-*T2A*-co *luxC*) (DEC(2A)). In addition, co *luxB* was fused with co *Frp* using GGGGS (G4S) (24). As a result, the fusion of co *luxB* and co *Frp*, BF(G4S), increased the luminescence intensity in cells compared to that of co lux (*SI Appendix*, Fig. S5*B*). In contrast, DEC(2A) did not enhance the intensity, which was also similarly reported previously (23), Therefore, by introducing the Kozak sequence (25) downstream of the 2A peptide, DEC(KZK) (*SI Appendix*, Fig. S5*A*), the luminescence intensity was improved (*SI Appendix*, Fig. S5*B*). The optimized co lux operon-expressing plasmids, BF(G4S) and DEC(KZK), were co-transfected with NLXs plasmids into HEK293T cells. The luminescence of the introduced NLXs was confirmed by microscopy using an EM-CCD camera for single-cell imaging. It was also shown that each luminescence color could be separated using specific optical filters (*SI Appendix*, Fig. S6*A*). Subsequently, we performed multiplex auto-bioluminescence imaging by mixing HEK293T cells expressing CNLX, YNLX, and RNLX (Fig. 2*C*). As with the mixed *E. coli* results, the cells could be distinguished by their respective wavelengths using spectral unmixing. We also conducted multiplex auto-bioluminescence imaging of subcellular structures by fusing RNLX with histones (H2B-RNLX) and YNLX with the plasma membrane (Lyn-YNLX), expressing them in HEK293T cells (*SI Appendix*, Fig. S6*B*). The luminescence signals from H2B-RNLX and Lyn-YNLX were separated by optical filtering and successfully showed correct localization.

We discovered that these optimized constructs could be transiently expressed in plant hosts without being optimized for plant codon usage. As a result of introducing the co lux operon constructs into the leaves of *Nicotiana benthamiana*, auto-bioluminescence from the leaves was successfully detected (Fig. 2*D* and *SI Appendix*, Fig. S7*A*). It also showed that NLXs produced luminescence intensities comparable to those of fungal luciferase (nnLuz) (26) (*SI Appendix*, Fig. S7*B*). No significant differences were observed in the gene expression levels of NLXs and nnLuz (*SI Appendix*, Fig. S7*C*). Compared to nnLuz, which generates a single color, our NLXs, capable of producing multicolor auto-bioluminescence, serve as versatile reporter genes for plant research. Thus, the NLX-based Lux operon demonstrated multiplex auto-bioluminescence observations in various living organisms.

### Application of NLXs as gene expression

Luciferase-based probes such as *Renilla* luciferase (Rluc) and firefly luciferase (Fluc) (27) have been widely used as reporter genes. However, luciferase-based probes still rely on luciferin addition. Thus, the ability to monitor the dynamics of reporter genes at the single-cell level for long-term observation without a decrease in luminescence intensity is limited. To evaluate whether our NLXs can also be used in reporter gene assays, we carried out a Wnt-reporter assay utilizing the Wnt-responsive promoter 7×TCF(5). The addition of LiCl as an agonist chemical to the Wnt protein resulted in luminescence from the cell-encoded 7×TCF-YNLX plasmid (Fig. 3*A*). We also checked whether the addition of LiCl or Wnt-activation can affect the luminescence intensity of cells expressing Lux systems (co lux). No significant changes in auto-bioluminescence intensity were observed after the addition of LiCl compared to cells expressing the Wnt-responsive promoter 7×TCF (7×TCF-YNLX) (Fig. 3*B*). To track the dynamics of Wnt responsiveness at the single-cell level, we continued to express 7×TCF-YNLX in HEK293T cells for 16 h (*SI Appendix*, Fig. S8*A*). Bioluminescence was observed four hours into the observation period and continued to be expressed (*SI Appendix*, Fig. S8*A* and Movie S1). Thus, the reporter gene based on YNLX was successfully used to monitor the dynamics of the reporter gene for long-term imaging.

**Fig. 3.**
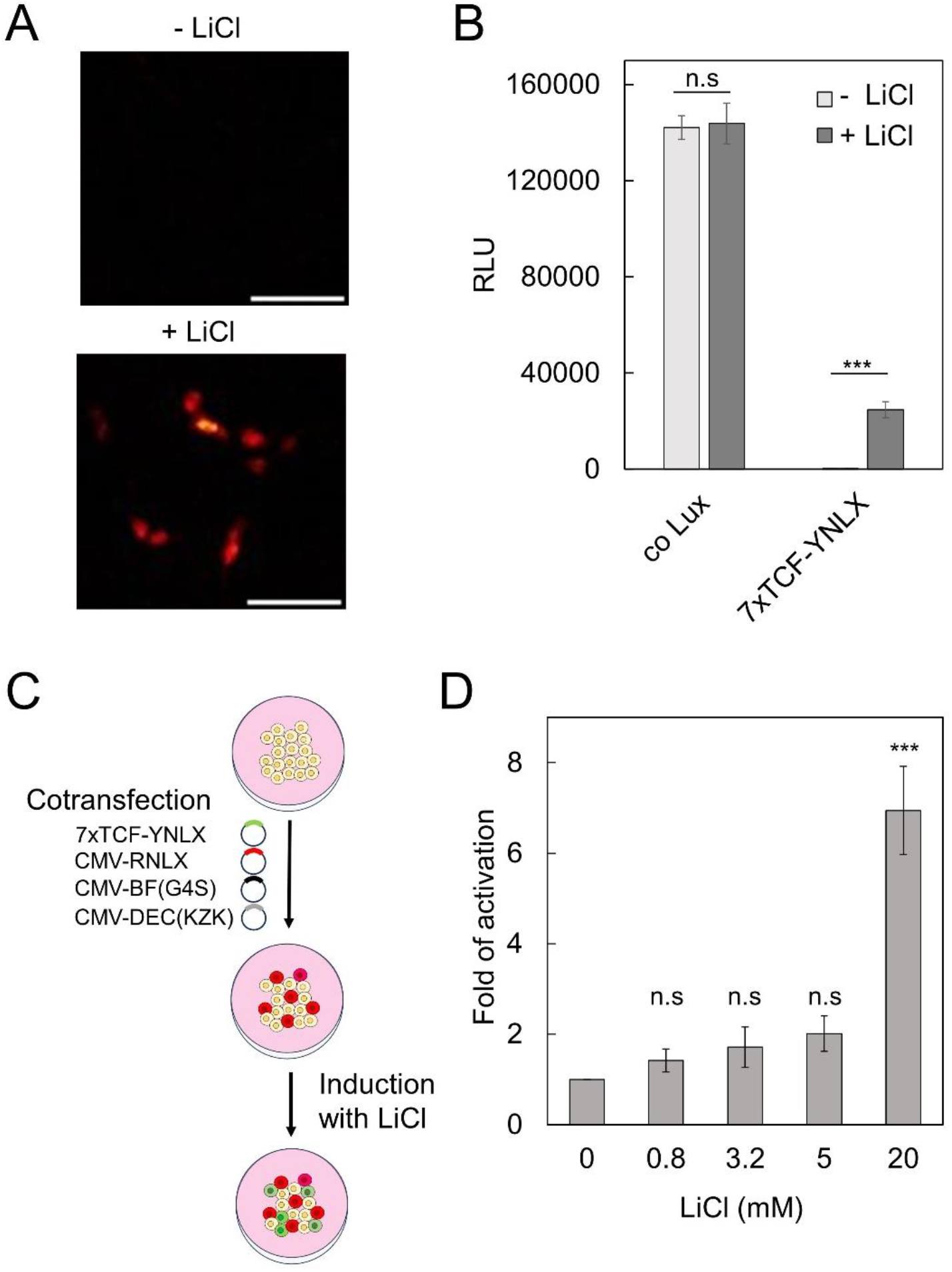
Applications of NLXs in gene expression. Reporter assay for Wnt gene expression by YNLX. Auto-bioluminescence imaging (*A*) and auto-bioluminescence intensity (*B*) of HEK293T cells expressing co lux and 7×TCF-YNLX were measured at 16h with or without the addition of 40 mM LiCl. Scale bars, 100 μm: ×20 magnification. RLU is the relative light unit. Data are mean ± s.d. *n* = 3, *** *p* < 0.001. (*C*) Schematic of the multiplex assay. HEK293T cells transiently transfected with 7×TCF-YNLX, CMV-RNLX, and auto-bioluminescent parts (CMV-BF(G4S) and CMV-DEC(KZK)) plasmids were treated with LiCl to induce Wnt expression. (*D*) Auto-bioluminescence intensities of 7×TCF-YNLX upon the addition of various concentrations of LiCl. Bioluminescence intensities were measured 16 h after the addition of various concentrations of LiCl. Data are mean ± s.d. *n* = 3, \*\*\**p* < 0.001.

We next demonstrated this reporter’s utility in transient transfection for high-throughput screening (HTS) by co-transfecting CMV-RNLX and 7×TCF-YNLX in HEK293T cells, producing normalized data accounting for varying transfection efficiencies (Figs. 3*C* and 3*D*). Unlike previous reporter assays utilizing Lux (28), which relied on Fluc or Rluc to normalize luminescence intensity and required cell lysis in downstream processes, our NLX system eliminates the need for additional luciferases and the requirement for cell lysis. This feature enhances convenience and cost-effectiveness, making it particularly advantageous for HTS applications.

### Application of NLXs as peptide-based indicator for ion and molecule

We expanded the application of Lux luciferase by developing a peptide-ligand-based indicator. To this end, we employed a ratiometric indicator strategy (Fig. 4*A*) to minimize signal drift due to changes in cell shape, focus drift, or variations in substrate consumption from auto-bioluminescence mechanisms. Consequently, we developed BRET-based indicators to detect crucial ligands such as calcium and ATP in living organisms. To develop a calcium indicator based on Lux, we fused the Ca^2+^-sensitive troponin-C peptide (TnC) (29) into the site between Venus and LuxA in the YNLX plasmid. First, we screened Venus variants and circularly permutated Venus variants (30) to achieve the highest dynamic range upon Ca^2+^ addition (*SI Appendix*, Fig. S9). Among them, cp173Venus-TnC-LuxA (YNLX(Ca^2+^)) showed the largest dynamic range (181%) compared with the other variants (Fig. 4*B*). The *K*_d_ value for Ca^2+^ in this construct was 570 nM and the Hill coefficient was 1.02 (Fig. 4*C*). Furthermore, we observed Ca^2+^ changes after the addition of ionomycin and 10 mM CaCl_2_ (Fig. 4*D*).

**Fig. 4.**
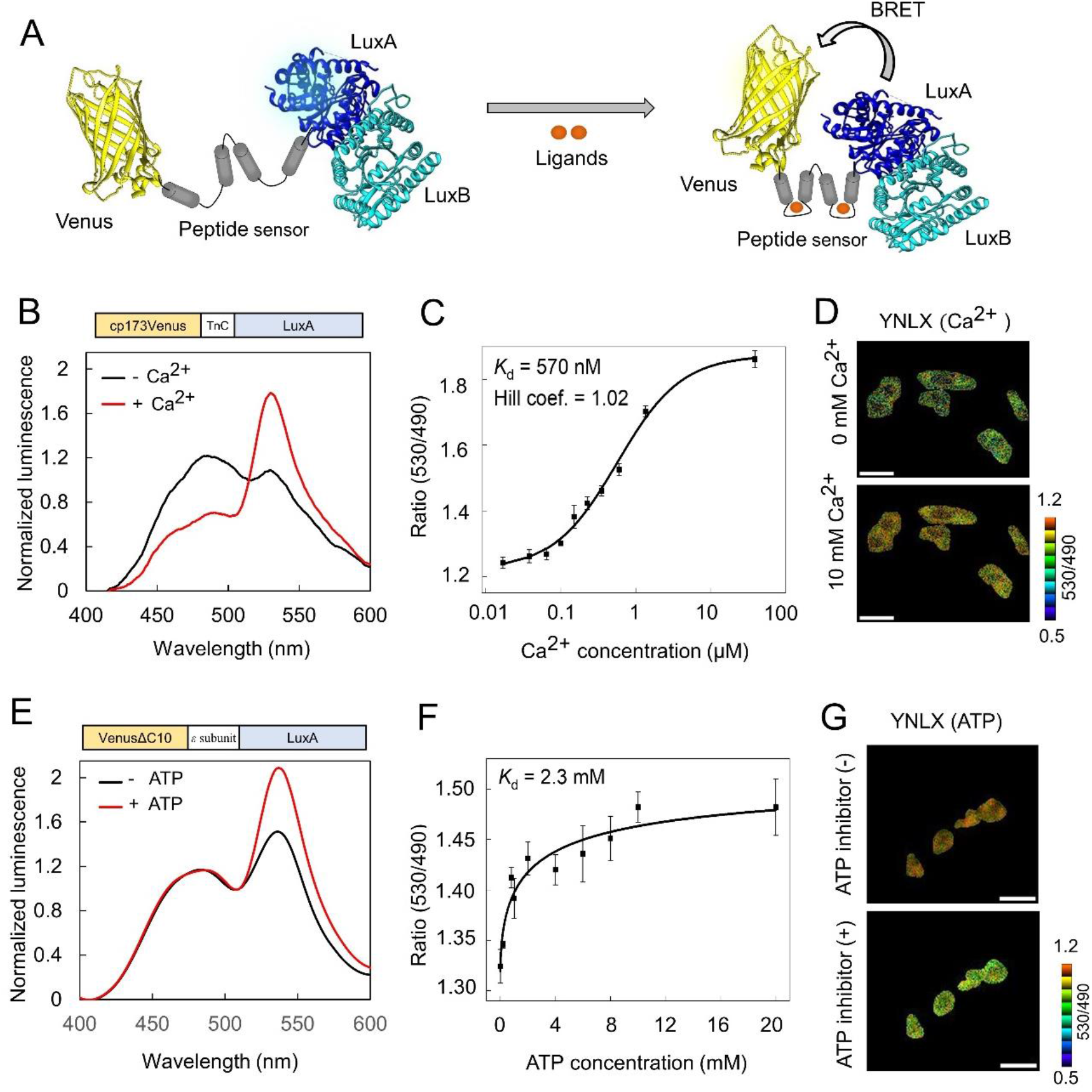
Applications of NLXs as peptide-based sensors. (*A*) Schematics of peptide-based sensor based on NLXs with ratiometric strategy. (*B*) Emission spectra of YNLX (Ca^2+^) with or without Ca^2+^. Bioluminescence intensities were normalized by the isoemission point. (*C*) Ca^2+^ titration curve of YNLX(Ca^2+^). The bioluminescence ratio was calculated from the peaks at 490 and 530 nm. (*D*) Auto-bioluminescence imaging of YNLX(Ca^2+^). BRET ratio (pseudocolor images) of HEK293T cells expressing YNLX(Ca^2+^) before and after addition of 10 μM ionomycin and 10 mM CaCl_2_. Scale bars, 50 μm: ×40 magnification. (*E*) Emission spectra of YNLX (ATP) with and without ATP. (*F*) ATP titration curve of YNLX(ATP). The bioluminescence ratio was calculated from the peaks at 490 and 530 nm. (*G*) Auto-bioluminescence imaging of YNLX (ATP). BRET ratio (pseudocolor images) of HEK293T cells expressing YNLX(Ca^2+^) before and after the addition of an ATP inhibitor (20 μg/mL oligomycin A and 20 mM 2-deoxyglucose). Scale bars, 50 μm: ×40 magnification.

To develop an ATP indicator based on Lux, we fused the ε subunit (31) into the site between Venus and LuxA in YNLX, resulting in Venus-ε subunit-LuxA (YNLX(ATP)). YNLX(ATP) showed elevated BRET efficiency in the presence of ATP, with the *K*_d_ value for ATP of this construct being 2.3 mM (Figs. 4*E* and 4*F*). We also successfully imaged intracellular ATP depletion upon addition of an inhibitor of ATP production (oligomycin and 2-deoxyglucose) (Fig. 4*G* and *SI Appendix*, Fig. S10). Therefore, YNLX was successfully applied as a peptide-based biosensor using a ratiometric strategy.

## Discussion

Autonomous bioluminescence technology facilitates time-lapse imaging of biological processes without the need for luciferin, which is advantageous as it eliminates the issue of luminescence decay. In addition, this technology is cost-effective for high-throughput screening, making it a cost-efficient option for luminescence imaging. However, the current state of auto-bioluminescent technology is primarily limited to observing individual biological phenomena, which hampers its utility for multiplexed autonomous imaging. Through the optimization of the LuxA position and BRET donor pairs, we successfully developed five color variants of auto-bioluminescence based on Lux luciferase. These color variants not only shifted the emission spectrum but also increased the luminescence intensity compared to that of the wild-type Lux. The cyan version, CNLX, is a brighter variant, potentially up to ten times brighter than the wild-type Lux, attributed to changes in quantum yield influenced by the BRET phenomenon. The use of NLXs has enabled us to perform multiplex auto-bioluminescent imaging, overcoming the limitations of wild-type Lux and other auto-bioluminescent applications such as fungal luciferase (Luz). A notable feature of NLXs is their ability to facilitate auto-bioluminescence in various organisms, including bacterial, mammalian, and plant hosts.

In comparison, the Luz system is only capable of fully establishing auto-bioluminescence in plant hosts, and not in other hosts, such as mammalian cells, because of the inability to synthesize caffeic acid as luciferin precursor in their basal metabolic pathway (3). Additionally, our system contradicts the conventional belief that Lux causes low light intensity in the plant host (32). In fact, our findings revealed that we were able to grow transiently multicolor bright plants with light intensity comparable to Luz. This makes our system an ideal tool for plant research. Furthermore, we have expanded auto-bioluminescent applications, traditionally limited to single sensors, by developing subcellular tags, multiple gene assays, and a proof-of-concept peptide-based sensor for auto-bioluminescent Ca^2+^ and ATP imaging in longer observations. We are also focused on the importance of red-shifted emission of RNLX for imaging animal tissues. This is crucial because it allows for distinction from the signal absorption of deep tissues, particularly from hemoglobin (33, 34). In addition, the yellow-shifted emission of YNLX is essential for plant research because of its absorption spectrum of chlorophyll (35). Furthermore, the improvement in the enzymatic catalytic efficiency of LuxA through an increase in the *k*_cat_ value has the potential to significantly elevate its luminescence intensity, bringing it to a level comparable to that of eNL (6). Overall, we believe that our NLXs could be a “game-changing” development in bioluminescent technology, offering multipurpose applications in the future.

## Materials and Methods

*SI Appendix, SI Materials and Methods* provided the details of the materials and methods used in this study, including of plasmid generation, luminescent characterization, cell culture, transient expression in plant leaves and quantitative analysis of gene expression. In brief, NLXs and its modified genes were cloned into pRSET_B,_ pCDNA3.1(+), and pRI201-AN for bacterial, mammalian, and plant expression, respectively. Bioluminescence imaging was performed with an inverted microscope based on the IXploreTM Live system equipped with EM-CCD camera.

Nucleotide sequences of NLX constructs are available as Datasets S1.

## Acknowledgments

We thank Prof. Stefan W. Hell for providing the cDNA for co lux. We also thanks to Dr. Kenji Osabe for the advice of luminescent proteins characterization. This work was partly supported by grants from the JST CREST program (No. JPMJCR20H9 to T.N.), the Ministry of Education, Culture, Sports, Science and Technology (MEXT) (No. 18H05410 to T.N.), Japan Society for the Promotion of Science (JSPS) (No. 22H00409 to T.N.), and MEXT scholarship to S.H.K.

