## Supplementary material for "Autonomous multicolor bioluminescence imaging in bacteria, mammalian, and plant hosts": 240429 NLX SUPPLEMENT Biorxiv.pdf

### SI Materials and Methods

**General methods.** DNA oligonucleotides were purchased from Hokkaido System Science. KOD-Plus-Neo (Toyobo Life Science) was used for the PCR amplification. PCR products were purified by agarose gel electrophoresis using a QIAEX II gel extraction kit (QIAGEN). *Dpn*I-treated was used to degrade the parental vector from the inverse-PCR vector. Small-scale plasmid DNA was obtained from a 1.5 mL LB-liquid bacterial culture by alkaline lysis and ethanol precipitation. Large-scale plasmid DNA was obtained from bacterial pellets from 200 mL of LB liquid culture by alkaline lysis, PEG-8000 precipitation, two phenol/chloroform extraction rounds, and isopropanol precipitation. DNA sequencing of the cDNA constructs was performed using the BigDye Terminator v1.1 Cycle Sequencing kit (Life Technologies). Other reagents such as FMN and BNAH were purchased from Tokyo Chemical Industry.

**Construction of multicolor Lux.** The *luxAB* operon in the $pRSET_B$ vector was used as the starting material to develop multicolor Lux for *E. coli* expression. cDNA of all fluorescent proteins (FPs) from $pRSET_B$ encoding mTQ2, sfGFP, Venus, mKOκ, mScarlet, mScarlet-I, CyOFP, and mCherry-XL were amplified by PCR and subcloned into the N-terminus of luxA in $pRSET_B$-*luxAB* with the addition of EL (glutamic acid-leucine) linkers using the TEDA method (1). The cDNAs of C-terminally deleted FPs mutants (FPsΔC10-*luxAB*), except mKOκ, were amplified by inverse PCR and circularized using the TEDA method. The C-terminally deleted FPs were also subcloned into the N- or C-terminus of LuxB to generate BRET-based LuxB using the TEDA method.

**Construction of mammalian expression vectors.** For mammalian expression, human codon-optimized lux from co lux (2) was used to generate auto-bioluminescent mammalian cells. The cDNA of FPs in NLXs were subcloned into the N-terminus of co luxA in the pcDNA3.1(+) vector using the TEDA method. The fusion of co luxB with co Frp was fused by using the GGGGS linker using the TEDA method. For the 2A peptide, P2A and T2A were fused downstream of co luxD and co luxE, respectively, to yield co *luxD-P2A*-co *luxE-T2A* in the pcDNA3.1(+) vector, using the TEDA method. Subsequently, luxC was fused downstream of T2A from co luxD-P2A-co luxE-T2A to yield co luxD-P2A-co luxE-T2A-co luxC. For nucleus localization, we subcloned the PCR-amplified of H2B from pcDNA3-ReNL-H2B into the N-terminus of RNLX in pcDNA3.1(+) vectors. For plasma membrane localization, we replaced YeNL from pcDNA3-lyn-YeNL with YNLX, using the TEDA method.

**Construction of plant expression vectors.** For plant expression, we subcloned NLXs-P2A-co luxBFrp and co luxD-P2A-co luxE-T2A-co luxC (DEC(KZK)) from pcDNA.1(+) into pRI201-AN (Takara Bio), which was linearized using *Sac*I and *Nde*I in MCS1 from the vector. The PCR-amplified NLXs-P2A-co luxBFrp and co luxD-P2A-co luxE-T2A-co luxC fragments, which have overlapping regions with the pRI201-AN vector (in the *Sac*I and *Nde*I sites), were ligated using the TEDA method to yield NLXs-P2A- co luxBFrp/pRI201-AN and co luxD-P2A-co luxE-T2A-co luxC/pRI201-AN. To construct a single plasmid vector for plant expression, co luxD-P2A-co luxE-T2A-co luxC/pRI201-AN was subcloned into MCS2 of NLXs-P2A- co luxBFrp/pRI201-AN using the TEDA method.

**Construction of gene expression reporter genes, $Ca^{2+}$, and ATP indicators.** Wnt-responsive enhancer 7×TCF and minimal CMV promoter were PCR-amplified from pT2-7×TCF-NLS-YNL (3) and subcloned into pcDNA3.1(+)-YNLX to obtain pcDNA3.1(+)-7×TCF-minCMV-YNLX. For $Ca^{2+}$ indicators, Troponin C (TnC) (4) was PCR-amplified from $pRSET_B$-Twitch-2B and subcloned into the N terminus of LuxA in $pRSET_B$-YNLX. A series of BRET-based $Ca^{2+}$ indicators were constructed by replacing VenusΔC10 in $pRSET_B$-YNLX with circularly permutated Venus (cp173Venus-TnC-LuxA) variants from Yellow cameleon (YCs) variants. Next, cp173Venus-TnC-LuxA was subcloned into pcDNA3.1(+)-YNLX vector for mammalian expression. For the ATP indicator, ε subunit (5) was PCR-amplified from $pRSET_B$-AT1.03, subcloned into the N-terminus of LuxA of $pRSET_B$-YNLX, and pcDNA3.1(+)-YNLX. All constructs were sub-cloned using the TEDA method.

**Protein expression and purification.** NLX proteins and those modified with an N-terminal polyhistidine tag were expressed in *E. coli* JM109 (DE3) at 23 °C for 60 h in liquid LB medium. The cells were collected by centrifugation and ruptured using a French press (ThermoFisher Scientific). The supernatant was purified using Ni-NTA agarose affinity columns (Qiagen), washed with 10 mM imidazole, and eluted with 100 mM imidazole. Finally, the buffer elution was changed to 20 mM HEPES (pH 7.4) using a PD-10 column (GE Healthcare).

**Protein characterization.** The luminescence characterization of recombinant proteins was measured using the BNAH method (6). Luminescence intensity was measured from 0.8 µM of luminescent proteins and mixed with 10 µM FMN, 20 µM decanal, and 100 µM BNAH (in auto-dispenser) using a microplate reader (SH-9000, Corona Electric) with 1 s exposure. The emission spectra were measured from 8 µM luminescent proteins of NLXs mixed with 10 µM FMN, 20 µM decanal, and 100 µM BNAH using a photonic multichannel analyzer PMA-12 (Hamamatsu Photonics) with 20 s exposure. The luminescent quantum yields were measured from the total light output by the complete consumption of 20 nm decanal using a microplate reader (SH-9000, Corona Electric) with 0.1 s exposures. The photon count of the microplate reader detector was determined using the luminol photon calibration method as reported previously (7). The final concentrations of proteins, FMN, and BNAH were 200 nM, 10 µM, and 100 µM BNAH (in an auto-dispenser). For kinetic parameters, 200 nM of proteins with final decanal concentrations of 0.01, 0.1, 0.5, 1, 5, 10, 50, and 80 µM were used. The initial reaction velocities were measured as the luminescence intensities for the initial 10 s and fitted to the Michaelis-Menten equation using Origin8 (OriginLab) software to estimate Michaelis-Menten constants ($K_m$) and maximum reaction velocities ($V_{max}$). *The* $k_{cat}$ values were calculated by dividing $V_{max}$ by the quantum yield and number of luciferase molecules. All experiments were performed at 37$^o$C and performed in triplicate. The averaged data were used for further analyses.

**Characterization of $Ca^{2+}$ indicator based on NLXs.** The emission spectra of the purified $Ca^{2+}$ indicator based on Lux and its variants were measured using PMA-12 (Hamamatsu Photonics) with 20 s exposure. The final protein concentration was 8 µM and then mixed with 10 µM FMN, 20 µM decanal, and 100 µM BNAH. $Ca^{2+}$ titrations were performed by the reciprocal dilution of $Ca^{2+}$ -free and $Ca^{2+}$-saturated buffers containing 10 mM MOPS, 100 mM KCl, and 10 mM EGTA with or without 10 mM $Ca^{2+}$ added as $CaCO_3$, at pH 7.2, 25 °C. The free $Ca^{2+}$ concentrations were calculated using 0.15 mM as the apparent $K_d$ value of EGTA for $Ca^{2+}$. The $Ca^{2+}$ titration curve was used to calculate the apparent $K_d$ value using nonlinear regression analysis. The averaged data were fitted to a single Hill equation using Origin8 software (OriginLab).

**Characterization of ATP indicator based on NLXs.** The emission spectra of the purified ATP indicator based on Lux were measured at 25 ˚C in HEPES using a microplate reader (SH-9000, Corona Electric) with 1s exposures. The final protein concentration was 8 µM and then mixed with 10 µM FMN, 20 µM decanal, and 100 µM BNAH. The final ATP solution (10 mM) was mixed with equimolar $MgCl_2$ to obtain the MgATP complex.

**Mammalian cell culture and transfection.** HEK293T cells were grown in Dulbecco's Modified Eagle's Medium (DMEM) (Sigma-Aldrich) supplemented with 10% fetal bovine serum (FBS) at 37 °C in 5% $CO_2$. The cells were cultured on collagen-coated 35 mm glass-bottom dishes or 12-well dishes. Transfection of recombinant DNA was performed using polyethylene imine (PEI Max 40 K; Polyscience), according to the manufacturer's instructions. The final concentration of all plasmids transfected was one µg/mL. In the case of six series of co lux, the total plasmids of co-transfected follow from the previous report (2) with a mixture of 83 ng co luxA, 83 ng co luxB, 83 ng co Frp, 250 ng co luxC, 250 ng co luxD, and 250 ng co luxE plasmids. The cells were then washed with phenol red−free DMEM/F12 and used for imaging or assay.

**Transient expression in *Nicotiana benthamiana* leaves by *Agrobacterium* infiltration.** Plant expression plasmids were transformed into *Agrobacterium tumefaciens* (GV3101) and cultured overnight on a shaker at 28 ˚C in LB medium under kanamycin selection. Bacterial cultures were collected and resuspended in infiltration buffer (10 mM $MgCl_2$, 10 mM MES pH 5.6, and 200 µM acetosyringone) to $OD_{500}$ = 0.6. The suspension was kept at room temperature in the dark for 2 h. The suspension was infiltrated using a needleless syringe to the abaxial side of *the N. benthamiana* leaves (5-6 weeks old). Three days after infiltration, bioluminescence was observed using a DSLR-based camera (SONY α7s, ISO 20000) with 60 s of exposure and a CCD-based camera (Vilber Fusion FX) with 1-60 s of exposure.

**Bioluminescence imaging.** For the luminescence image of NLXs expressed in *E. coli*, recombinant *E. coli* in the LB plates were exposed to SONY α7s, ISO 20000, for 10s of exposure. Auto-bioluminescence images of mixed *E. coli* expressing of CNLX, YNLX, and RNLX were picked from recombinant *E. coli* in LB plates into 10 µL of $H_2O$. 0.3 µL of each recombinant *E. coli* was introduced into 35 mm glass bottom dishes using the agar pad method. Luminescence imaging was acquired with an inverted microscope based on the IXplore™ Live system equipped with an EM-CCD camera (Andor iXon Ultra 888) with 8 min of exposure, EM-gain of 1000×, ×100 objective lens and 2 × 2 binning settings. Three filters, Olympus U-FCFP (460-510), Olympus U-FYFP (515-560), and Olympus U-FMCHE (600-690), were used to separate *E. coli* images.

Luminescence imaging of auto-bioluminescent NLXs expressed in HEK293T cells was acquired with an inverted microscope based on the IXplore™ Live system equipped with an x40 objective lens (Olympus, UPlanSApo, numerical aperture 1.4) was used. The emission signals were detected by an EM-CCD camera (Andor iXon Ultra 888) with 60s of exposure. Five filters, consist of Olympus U- Olympus FCFP (460-510), Semrock FF01-514/30 (510-550), Olympus U-FYFP (515-560), Semrock FF01-562/40, and Olympus U-FMCHE (600-690), were used to separate NLXs images. For mixed HEK293T cells expressing CNLX, YNLX and RNLX, three filters consist of Olympus U-FCFP (460-510), Olympus U-FYFP (515-560), and Olympus U-FMCHE (600-690), were used to separate the mixed cell images. For nucleus and plasma membrane localization tag imaging, HEK293T cells expressing H2B-RNLX and Lyn-YNLX were acquired with Olympus U-FYFP and Olympus U-FMCHE filters with 3 min of exposure, EM-gain of 1000×, ×100 objective lens and 2 × 2 binning settings.

For the Wnt reporter, an Olympus U-FYFP filter with 2 min of exposure, EM-gain of 1000x, x20 objective lens, and 2 × 2 binning settings was used. For YNLX($Ca^{2+}$), Olympus U-FCFP and Olympus U-FYFP filters with 2 min of exposure, EM-gain of 1000×, ×40 objective lens, and 2 × 2 binning settings were used with or without the addition of 10 mM ionomycin and 10 mM $CaCl_2$. For YNLX(ATP), Olympus U-FCFP and Olympus U-FYFP filters with 3 min of exposure, EM-gain of 1000×, ×40 objective lens, and 2 × 2 binning settings were used with or without the addition of ATP inhibitors (20 µg/mL oligomycin A and 20 mM 2-deoxyglucose). All imaging conditions were performed at 37 °C, 5% $CO_2$ environment in a stage-top incubator, STX (TOKAI HIT). All bioluminescent images were analyzed using ImageJ software (8) and MetaMorph software.
Luciferase assay

For the luciferase assay of the Wnt reporter, original co lux, 7×TCF-minCMV-YNLX with or without co-transfected with CMV-RNLX plasmids were transfected into HEK293T cells on 12 well dishes. After 12 h of transfection, cells were treated with or without LiCl for 16 h. The cells were then transferred to a 96-well white microplate and luminescence was measured using a multimode plate reader (Spectra Max iD5, Molecular Devices) for 1 s of exposure. The luminescence intensities at 530 nm (YNLX signal) and 590 nm (RNLX signal) emission spectra were used to determine the effect of various LiCl concentrations. The reported unit was the luminescence intensity (RLU) of the 530 nm signal over the 590 nm signal. To derive fold of activation, all values were normalized to the corresponding non-treated control.

**Quantitative analysis of NLXs and nnLuz genes.** *N. benthamiana* leaves expressing NLXs and nnLuz were homogenized with metal beads, and total RNA was extracted using NucleoSpin® RNA Plant (Takara Bio). One microgram of total RNA was reverse transcribed to first-strand cDNA using ReverTra Ace™ qPCR RT Master Mix (Toyobo). Real-time quantitative RT-PCR was performed on a StepOne Real-Time PCR system (Thermo Fisher Scientific) using PowerUp SYBR Green Master Mix (Thermo Fisher Scientific). The baseline, threshold cycles ($C_T$), and comparative $C_T$ ($\Delta\Delta C_T$) were calculated using StepOne software v2.3. The result was normalized by *PP2A* (9) gene expression, and the relative gene expression was analyzed by comparative $C_T$ ($\Delta\Delta C_T$).

**Data analysis and statistical methods.** All bioluminescent images from the microscope were analyzed using ImageJ and MetaMorph software. The bioluminescence images of the microscope were subtracted from the background, and the cosmic rays were processed with an adaptive median filter without affecting the original brightness or morphology of the cells using ImageJ software. The pseudocolor images were used for microscopy images with “Red Hot” color for total luminescence intensity and a specific color for each cell according to specific wavelengths. The “16 color” was used to discriminate luminescence intensity of plant imaging. To produce ratio images of the $Ca^{2+}$ and ATP sensors, the ratio images were processed with MetaMorph, and LIMD pseudocolor was used to distinguish the ratio values.

Data fitting and statistical analysis were performed using Origin8 (OriginLab). Statistical analysis was performed using unpaired Student’s *t-test* for comparing two parameter sets, and one-way ANOVA followed by post hoc Tukey’s honestly significant difference test was used to compare more data sets.

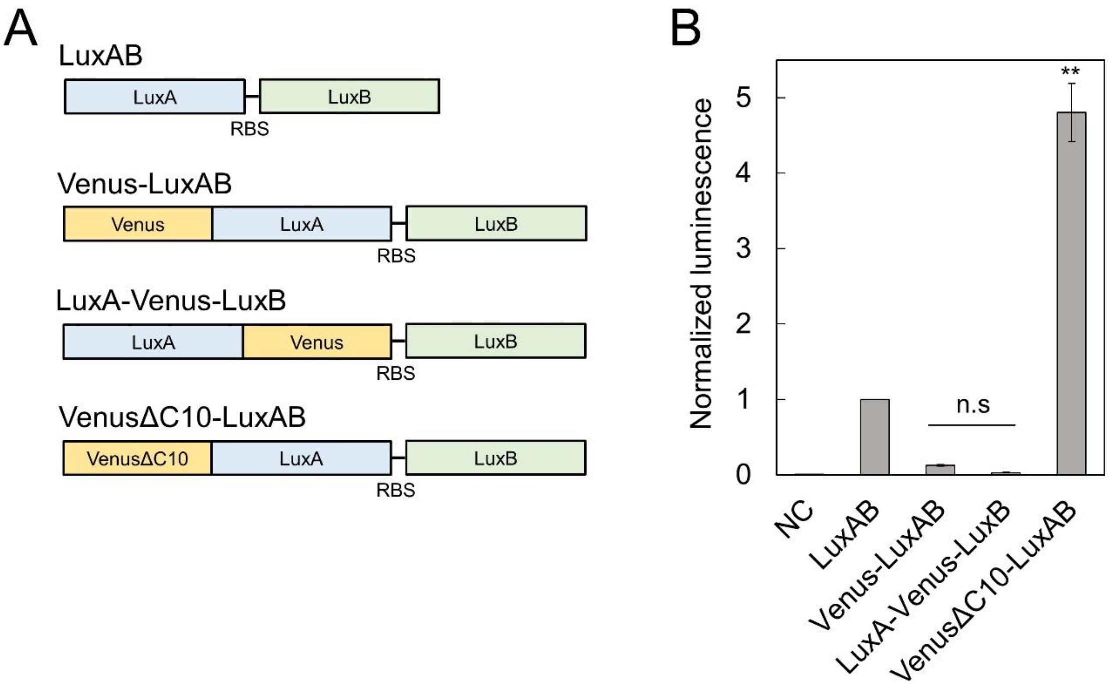


**Fig. S1.** Optimization of Venus fusion position to LuxA. (*A*) Schematics of Venus fusion position to LuxAB operon. RBS is ribosome binding site. (*B*) Bioluminescence intensity from equimolar amounts of with or without C-terminus deletion of Venus fused to LuxA. NC is HEPES solution without LPs. $n = 3$, $^{**}p < 0.05$.

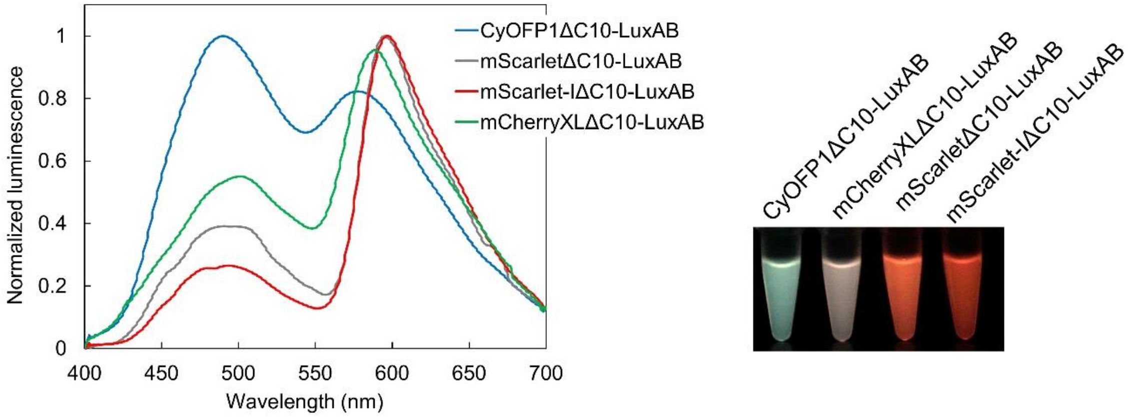


**Fig. S2.** Optimization of red variants Lux. Bioluminescence spectra of optimization for red variants of NLXs (left panel) and bioluminescence images of recombinant proteins (right panel). Bioluminescence intensities are normalized by peak intensity.

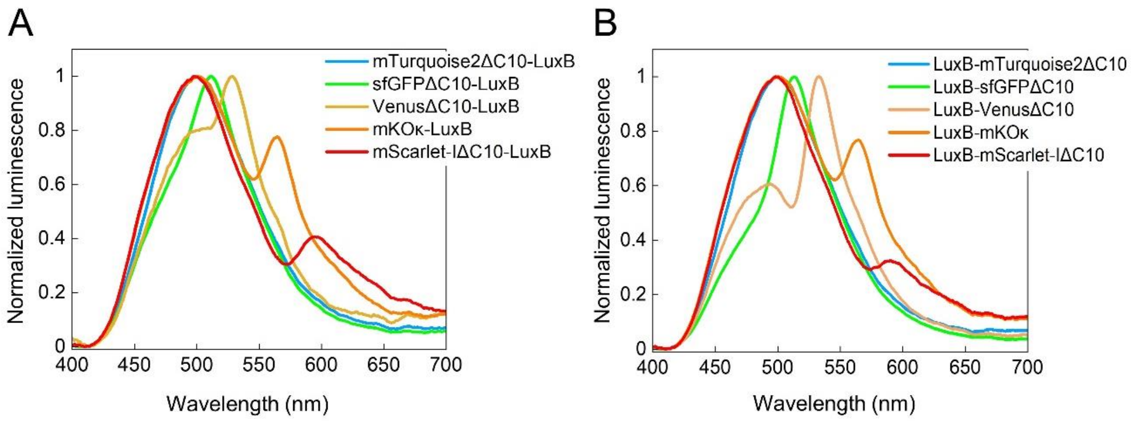


**Fig. S3.** Fluorescent proteins fusion to N or C-terminus of LuxB. Emission spectra of fluorescent proteins fused to N (*A*) or C-terminus (*B*) of LuxB. Bioluminescence intensities are normalized by peak intensity.

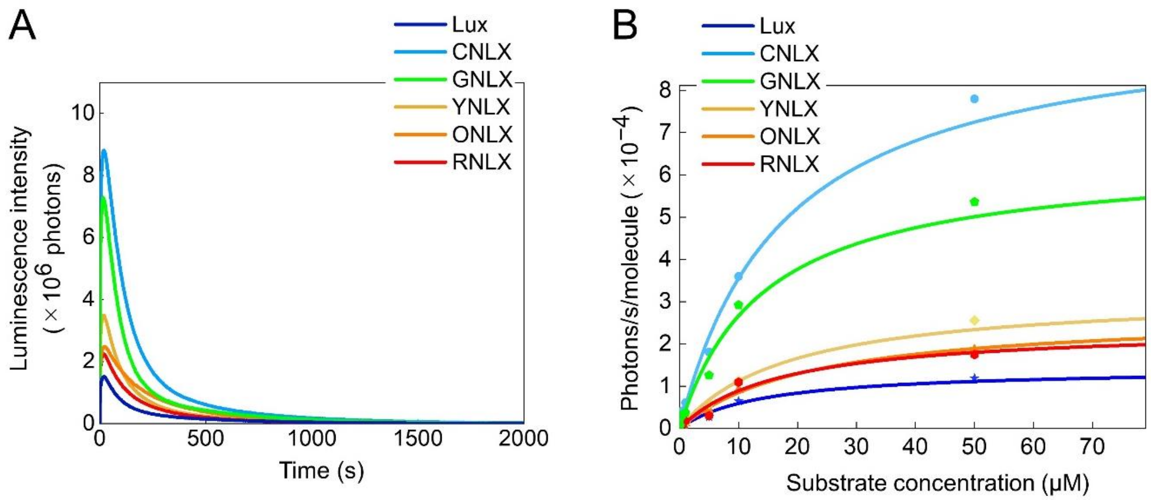


**Fig. S4.** LPs characterization. (*A*) Quantum yield (QY) measurement of NLXs. QY was estimated from the integrated light output until the reaction of substrate, decanal, approached completion. (*B*) Enzyme kinetics of NLXs. Kinetics parameters were estimated from the plot of the initial light output vs. the concentration of the decanal. The results are summarized in Table S2.

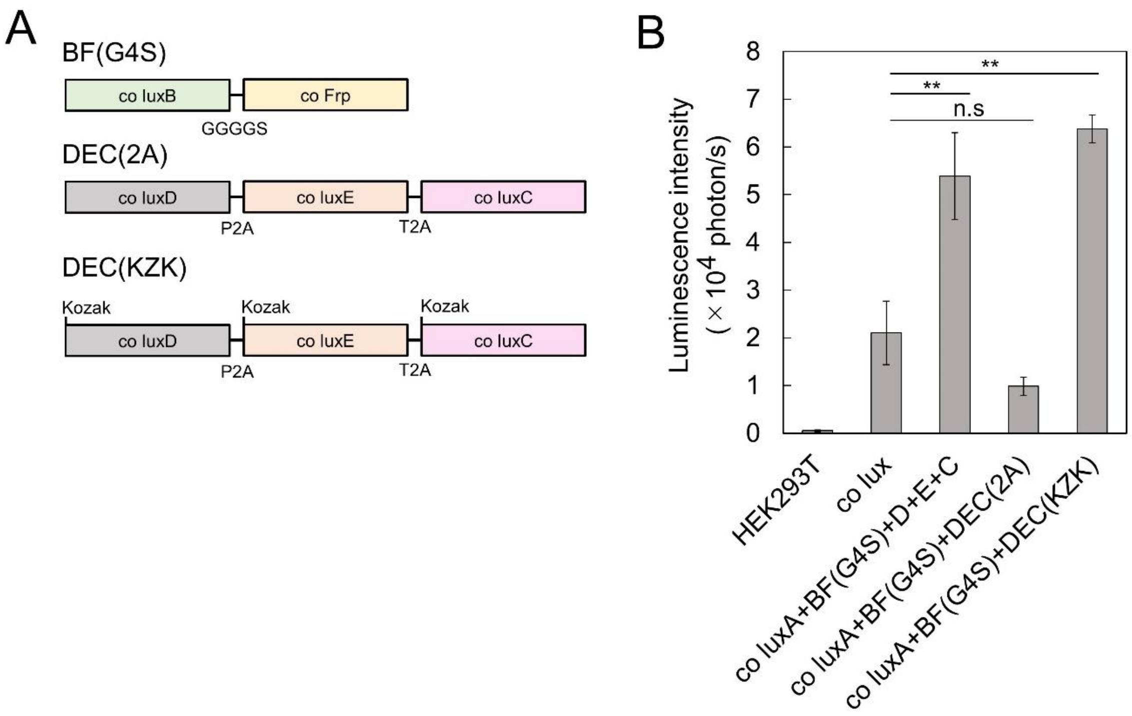


**Fig. S5.** Optimization of co Lux plasmids. (*A*) Schematics of the optimization co lux plasmids in pcDNA3.1(+) vector. (*C*) Total luminescence intensity of HEK293T cells only, HEK293T cells expressing of co lux plasmids (six plasmid co-transfected), five plasmids co-transfected by fusing co luxB with co Frp using GGGGS (G4S) linker, three plasmids transfected with co luxA, co luxBFrp, and co luxDEC (co luxD-P2A-co luxE-T2A-co luxC), and three plasmids transfected with co luxA, co luxBFrp, and co luxDEC with the addition of Kozak sequence in the downstream of 2A peptide. $n$ = 3, **$p$ < 0.05.

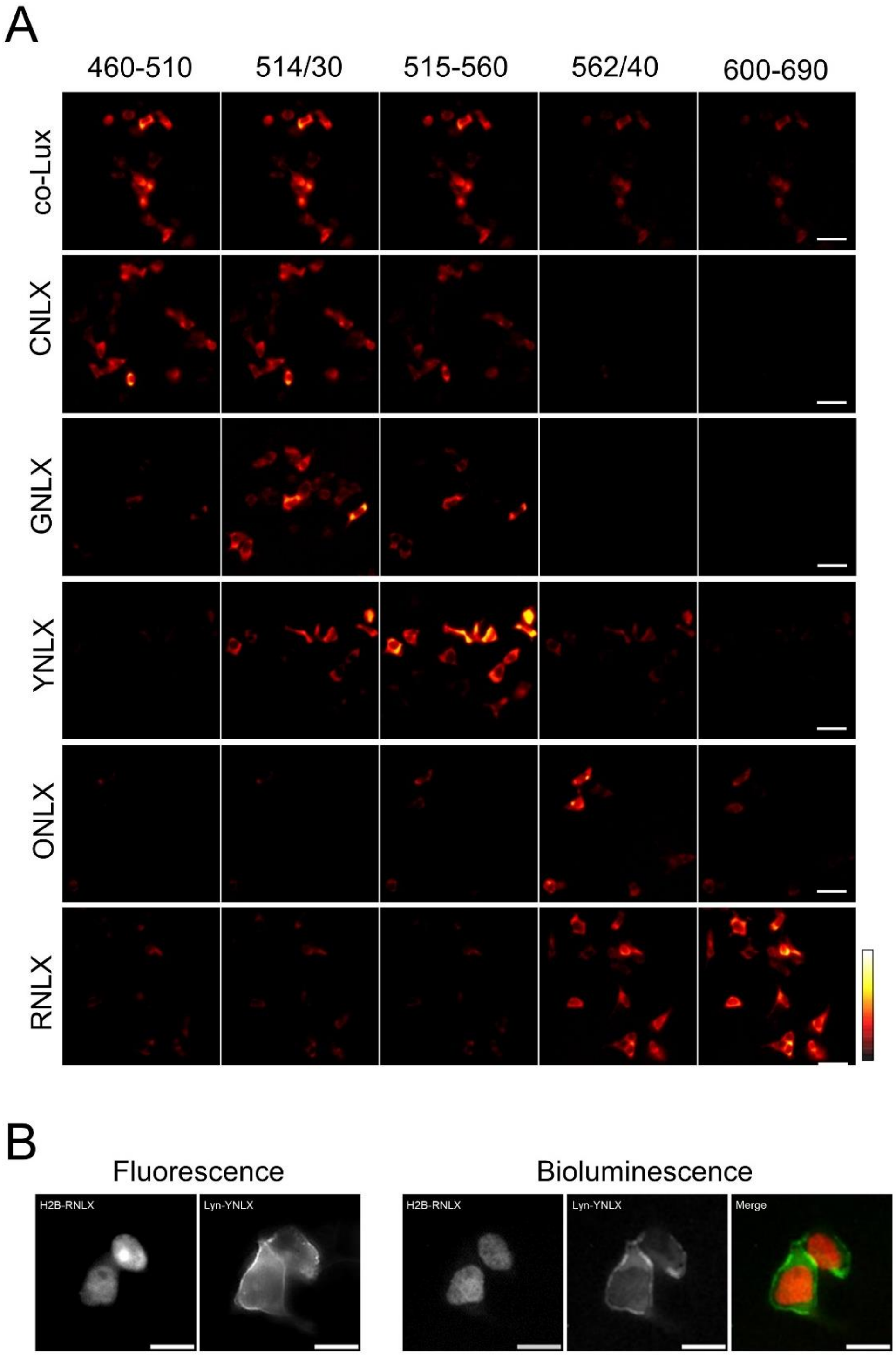


**Fig. S6.** Multiplexed imaging of single cells and subcellular localization tags in HEK293T cells. a, Auto-bioluminescence imaging of HEK293T cells expressing NLXs using different optical filters. Scale bars, 50 µm: ×40 magnification. b, Subcellular tags using NLXs. Bioluminescence imaging of HEK293T cells expressing NLXs targeted to the nucleus (H2B-RNLX) and plasma membrane (Lyn-YNLX). Scale bars, 20 µm: ×100 magnification.

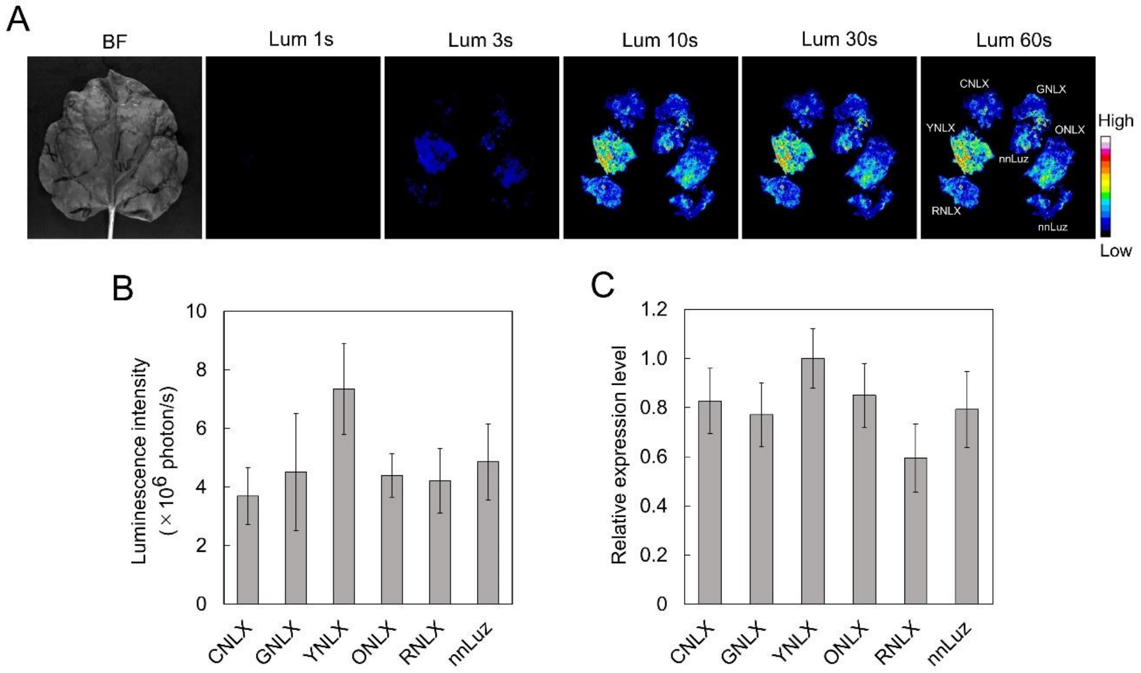


**Fig. S7.** Auto-bioluminescence of *Nicotiana benthamiana* leaves expressing NLXs genes and nnLuz by CCD-based camera. Auto-bioluminescence image (a) and total luminescence intensity (b) of the leaves expressing NLXs and nnLuz variants by *Agrobacterium* infiltration. Luminescence images were taken by CCD camera with several exposure time. Data were measured in triplicate, presented as luminescence intensity. c, Relative expression level of NLXs and nnLuz. The results were normalized to *the PP2A* gene expression.

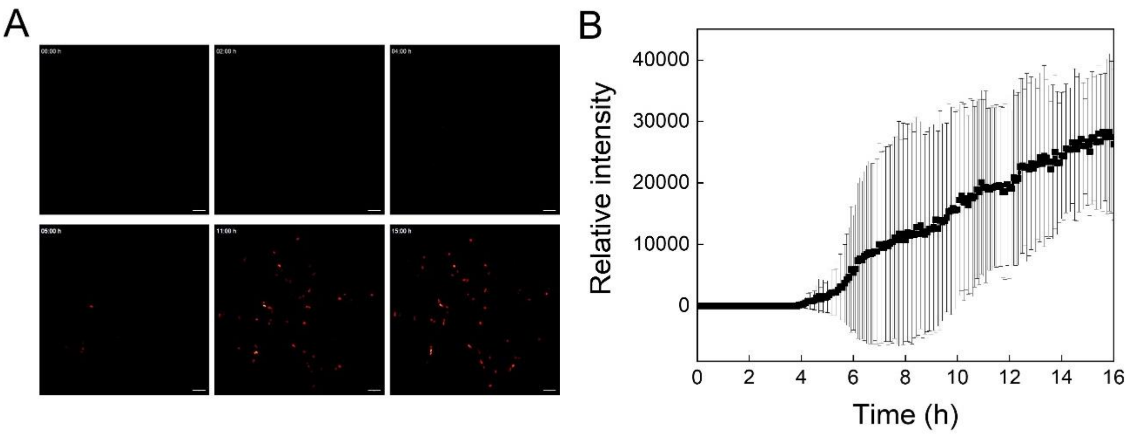


**Fig. S8.** Time-lapse imaging of 7×TCF-YNLX. Sequential images (a) and time course (b) of HEK293T cells expressing 7×TCF-YNLX upon the addition of 40 mM LiCl. Scale bars, 100 µm: ×20 magnification.

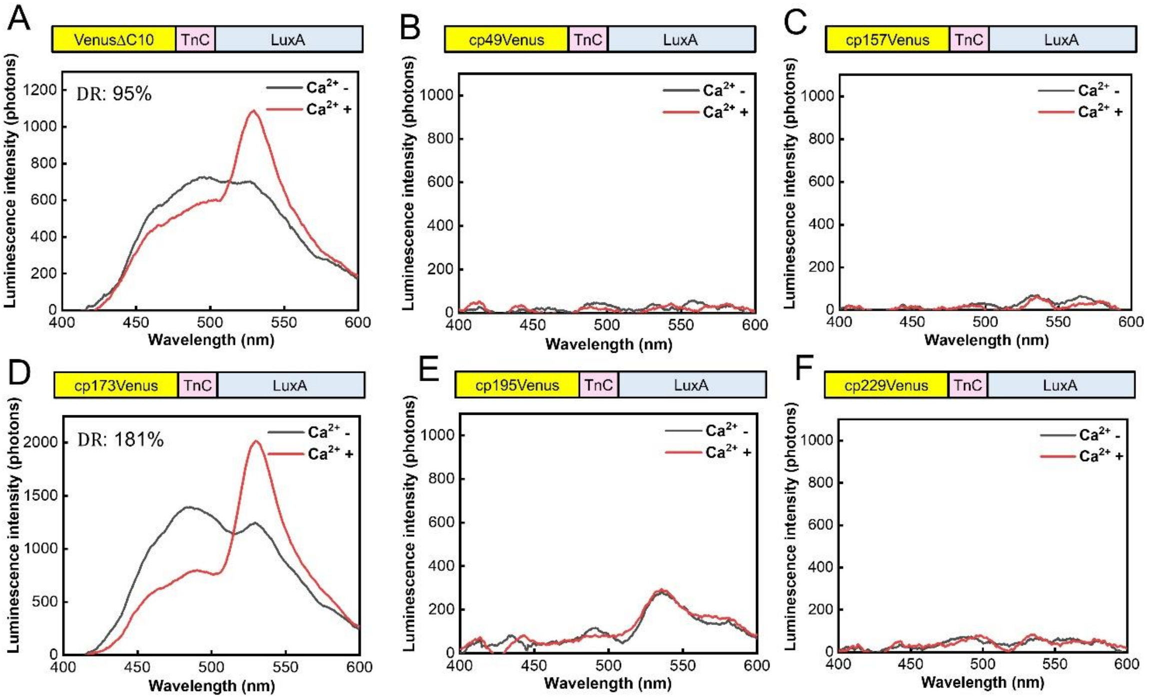


**Fig. S9.** Optimization of calcium indicators based on YNLX. Schematic (upper parts) and emission spectra (bottom parts) of calcium indicator based on NLX, a-f. Venus or cpVenus variants-TnC-LuxA, under 10 mM EGTA (black) and 20 µM $CaCl_2$ (red). Data were measured in triplicate, presented as luminescence intensity.

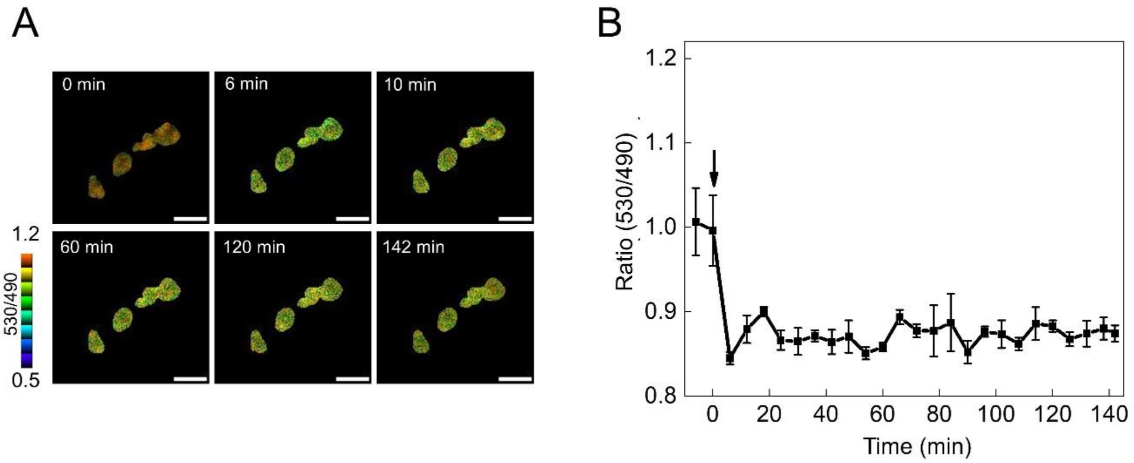


**Fig. S10.** Time-lapse BRET imaging of cytosolic ATP levels in living HEK293T cells using YNLX (ATP). a, Sequential images of the BRET ratio in HEK293T cells expressing YNLX(ATP). ATP depletion was monitored after addition of 20 µg/mL oligomycin A and 20 mM 2-deoxyglucose at time=0 (min). Scale bars, 50 µm: ×40 magnification. b, Time-course of the BRET ratio of HEK293T cells expressing YNLX(ATP). The arrow indicates the time of the addition of the inhibitors to the medium.

**Table S1.** Apparent BRET efficiencies of BRET Lux proteins.

| Name | $I_{acceptor}/I_{donor}$ |
|---|---|
| **LuxA-based BRET constructs** | |
| mTurquoise2ΔC10-LuxA (CNLX) | 1.06 |
| sfGFPΔC10-LuxA (GNLX) | 2.63 |
| VenusΔC10-LuxA (YNLX) | 7.69 |
| mKOκ-LuxA (ONLX) | 8.85 |
| mScarlet-IΔC10-LuxA (RNLX) | 3.85 |
| CyOFP1ΔC10-LuxA | 0.80 |
| mCherryXLΔC10-LuxA | 1.79 |
| mScarletΔC10-LuxA | 2.56 |
| **LuxB-based BRET constructs** | |
| mTurquoise2ΔC10-LuxB | 0.84 |
| sfGFP2ΔC10-LuxB | 1.33 |
| VenusΔC10-LuxB | 1.17 |
| mKOκ-LuxB | 0.68 |
| mScarlet-IΔC10-LuxB | 0.47 |
| LuxB-mTurquoise2ΔC10 | 0.82 |
| LuxB-sfGFP2ΔC10 | 1.68 |
| LuxB-VenusΔC10 | 1.54 |
| LuxB-mKOκ | 0.85 |
| LuxB-mScarlet-IΔC10 | 0.49 |

Comparison of BRET efficiency of BRET Lux proteins shown in Fig. 1 and Figs. S2 and S3. The BRET efficiency was measured by the ratio of luminescence intensities of the acceptor peak ($I_{acceptor}$) to that of the donor peak ($I_{donor}$).

**Table S2.** Enzymatic characteristics of luminescent proteins.

| Name | LQY | $K_m$ (μM) | $V_{max}$ (photon $s^{-1}$ $molecule^{-1}$) | $k_{cat}$ ($s^{-1}$) |
|---|---|---|---|---|
| Lux | 0.11 ± 0.01 | 30.48 ± 15.10 | $1.47 \times 10^{-4} \pm 1.72 \times 10^{-5}$ | $1.24 \times 10^{-3} \pm 1.45 \times 10^{-4}$ |
| CNLX | 0.64 ± 0.05 | 23.46 ± 7.74 | $9.43 \times 10^{-4} \pm 3.48 \times 10^{-5}$ | $1.58 \times 10^{-3} \pm 1.32 \times 10^{-4}$ |
| GNLX | 0.45 ± 0.03 | 24.58 ± 8.21 | $6.61 \times 10^{-4} \pm 7.74 \times 10^{-5}$ | $1.47 \times 10^{-3} \pm 1.72 \times 10^{-4}$ |
| YNLX | 0.27 ± 0.01 | 25.74 ± 6.74 | $3.66 \times 10^{-4} \pm 3.99 \times 10^{-5}$ | $1.35 \times 10^{-3} \pm 1.47 \times 10^{-4}$ |
| ONLX | 0.23 ± 0.01 | 40.91 ± 17.25 | $3.49 \times 10^{-4} \pm 1.31 \times 10^{-4}$ | $1.49 \times 10^{-3} \pm 5.60 \times 10^{-4}$ |
| RNLX | 0.18 ± 0.02 | 31.42 ± 12.61 | $3.64 \times 10^{-4} \pm 1.16 \times 10^{-4}$ | $1.82 \times 10^{-3} \pm 3.95 \times 10^{-4}$ |

LQY, luminescent quantum yield. Data are presented as mean ± S.D., $n$ = 3.

**Movie S1 (separate file).** Time-lapse imaging of 7×TCF-YNLX. Time lapse imaging of HEK293T cells expressing 7×TCF-YNLX upon the addition of 40 mM LiCl. Scale bar = 50 µm.

**Datasets S1 (separate file).** Nucleotide sequences of NLX constructs used in this study.
